# Protein language models reveal evolutionary constraints on synonymous codon choice

**DOI:** 10.1101/2025.08.05.668603

**Authors:** Helen Sakharova, Liana F. Lareau

## Abstract

Evolution has shaped the genetic code, with subtle pressures leading to preferences for some synonymous codons over others. Codons are translated at different speeds by the ribosome, imposing constraints on codon choice related to the process of translation. The structure and function of a protein may impose pressure to translate the associated mRNA at a particular speed in order to enable proper protein production, but the molecular basis and scope of these evolutionary constraints have remained elusive. Here, we show that information about codon constraints can be extracted from protein sequence alone. We leverage a protein language model to predict codon choice from amino acid sequence, combining implicit information about position and protein structure to learn subtle but generalizable constraints on codon choice in yeast. In parallel, we conduct a genome-wide screen of thousands of synonymous codon substitutions in endogenous loci in yeast, reliably identifying a small set of several hundred synonymous variants that increase or decrease fitness while showing that most positions have no measurable effect on growth. Our results suggest that cotranslational localization and translational accuracy, more than cotranslational protein folding, are major drivers of selective pressure on codon choice in eukaryotes. By considering both the small but wide-reaching effects of codon choice that can be learned from evolution and the strong but highly specific effects determined via experiment, we expose unappreciated biological constraints on codon choice.

## Introduction

Codon choice represents a fundamental puzzle in how biological information is encoded in genomes. The redundancy of the genetic code provides many ways to encode the same protein, but synonymous codons are not perfectly interchangeable. While the rate of synonymous substitutions is often used as a proxy for neutral evolution, genomes show biases towards some codons over others, and nominally silent mutations can have real consequences on function. These consequences are brought into relief by the growing importance of synthetic mRNAs for vaccines and therapeutics. Understanding the biological impact of codon choice is of deep practical concern for sequence design and genome interpretation.

Codon choice has a direct impact on the process of translation itself by changing the speed of translation elongation. Preferred codons are matched by abundant tRNAs, enabling faster and more accurate decoding (Ikemura, 1981; Mordret *et al*., 2019; Sun and Zhang, 2022; Drummond and Wilke, 2008; Kramer *et al*., 2010). Fast translation elongation generally leads to higher protein output, while slow translation elongation decreases output by triggering mRNA degradation and limiting translation initiation (Presnyak *et al*., 2015; Barrington *et al*., 2023; Lyons *et al*., 2024). Proteins that are in high demand are often encoded by fast codons, particularly in organisms like yeast with rapid growth rates that demand efficient resource utilization (Harrison and Charlesworth, 2011; de Oliveira *et al*., 2021; Bénitière *et al*., 2025). Our previous work showed that a neural network could maximize production of a simple fluorescent protein simply by minimizing total elongation time (Tunney *et al*., 2018). These observations might appear to imply that fast codons would be universally preferred.

However, simple rules for optimization ignore crucial yet idiosyncratic roles for slow codons arising from translational and cotranslational processes. Despite substantial experimental effort, the nature and impact of these processes is subtle and hard to predict, and we are far from a generalized understanding of when and where codon choice matters. Slow translation at the start of a gene is thought to help space out ribosomes and avoid ribosome collisions that trigger RNA decay (Tuller *et al*., 2010). Regions of slow translation may play an important role in cotranslational protein folding, allowing more time for earlier segments of the protein to fold before later segments are generated (Zhang *et al*., 2009; Kim *et al*., 2015; O’Brien *et al*., 2012). Similarly, slow translation can aid cotranslational localization by providing extra time for localization machinery to bind to an extruded peptide before the rest of the protein is produced (Pechmann *et al*., 2014; Mahlab and Linial, 2014; Acosta-Sampson *et al*., 2017).

The impact of translational and cotranslational processes on codon choice is further obscured by forces unrelated to translation. Nucleotide changes can add or disrupt mRNA structures, splice sites, or other important nucleotide motifs (Liu *et al*., 2021). Mutational biases and pressure to maintain a particular GC content add constraints unrelated to the function of the sequence *per se* (Sved and Bird, 1990; Sejour *et al*., 2023; Palazzo and Kang, 2021; Kudla *et al*., 2006). These layers combine into a complex landscape of subtle constraints shaping coding sequences. Nonetheless, the process of translation itself is central to the question of synonymous codon choice, and disentangling its direct impact remains a major challenge in understanding the genome. Our goal is to pinpoint where slow translation or fast translation are specifically required to produce functional protein and what mechanisms drive this requirement.

To uncover the constraints on codon choice imposed by translation, we turned to deep learning models that can capture subtle patterns in biological sequence. Evolution has conducted millions of years of experiments to determine what synonymous mutations are permissible, and recent advances in machine learning raise the possibility of learning generalizable rules of codon choice from this evolutionary record. The breakthroughs that enabled large language models have recently been extended to the language of biological sequence, creating foundation models such as the Evolutionary Scale Model (ESM2) that learn rich representations of protein sequence context by observing patterns across the full tree of life (Rives *et al*., 2021; Lin *et al*., 2023). The internal representations learned by these models capture substantial information about protein structure and function solely from amino acid sequence. If protein structure and function determine the translation speed profile necessary for proper cotranslational protein folding and localization, we posit that a foundation model like ESM2, trained only on protein sequence, would also intrinsically capture subtle yet detectable information about codon choice. Recent work applying machine learning to understand codon choice has focused on codon-level foundation models (Outeiral and Deane, 2024; Li *et al*., 2024; Constant *et al*., 2023) or amino acid to codon sequence encoder-decoder frameworks (Faizi *et al*., 2025; Sidi *et al*., 2025). These models recover many aspects of sequence constraints, but their predictions capture overall preferences for GC content or gene-level preferences for faster codons that can obscure specific constraints on translation. By employing a model that considers only protein sequence and ignores biases in codon composition between genes, we instead isolate translation-specific pressure on codon choice.

Here, we combine a protein language model and a genome-wide CRISPR editing screen to reveal the landscape of synonymous codon constraints in yeast. Our model predicts codon choice from amino acid sequence alone, combining context from a foundation model of protein sequences across the tree of life with information about codon choice spanning a set of distantly related budding yeast. We pair our model with a careful CRISPR retron editing screen to measure the fitness effects of thousands of synonymous mutations in endogenous loci across the yeast genome. In contrast to recent results (Shen *et al*., 2022), we find that synonymous mutations at a small but important subset of codon positions have verifiable fitness effects, while the majority of synonymous changes do not affect fitness. Our model and our experiments align to highlight cotranslational localization and translation accuracy, more than cotranslational folding, as forces shaping codon choice across evolution.

## Results

### A deep learning model to predict codon choice from protein sequence

Our overall goal is to learn contexts in which the choice of synonymous codon is important for function. There are two major challenges to this goal. First, the selective pressure on codon choice is real but generally weak (Hershberg and Petrov, 2008; Smith and Eyre-Walker, 2001). Most single synonymous substitutions are not expected to have measurable phenotypes, and most synonymous positions are under close to neutral selection, with little evolutionary signature. Second, nucleotide sequences are shaped by many forces. Mutational biases, selection for GC content, and functional constraints from binding sites and RNA structure exist alongside evolutionary pressures on the actual process of translation. These considerations led us to an approach that attempts to strip out these confounders to learn where codon choice *per se* is biologically important. We chose to employ a model that is trained only on amino acid sequences, not nucleotide sequences, to learn a rich embedding representing the structural and functional context at each position in a protein. To focus on the translation-related consequences of codon choice, this embedding would then be used to predict the translation elongation speed of the codon at a given position, rather than identity of the codon. To capture the complexity of eukaryotic translation without extensive sequence constraints from alternative splicing and other processes, we built our model and conducted our experiments on coding regions in yeast, which are relatively free of these additional constraints. Finally, and importantly, we put careful thought into training the model, avoiding simple memorization of conserved positions and gene-specific biases in codon choice. The result is a model that successfully identifies subtle and generalizable constraints on synonymous codon choice.

### Model built from protein sequence accurately predicts codon preferences

To build our model, we began by establishing labels of ‘fast’ or ‘slow’ for each codon in yeast based on translation elongation rates. Our labels were based on average transcriptome-wide elongation rates determined by ribosome profiling experiments in *S. cerevisiae*, providing an empirical measure in contrast to commonly used indirect measures such as codon frequency. To ensure that our classification was robust, we excluded codons with intermediate elongation rates as well as codons whose elongation rates were expected to vary between yeast species. This left a subset of 26 codons, corresponding to eight amino acids, labeled in a binary manner as either fast or slow (Figure 1A).

**Figure 1:**
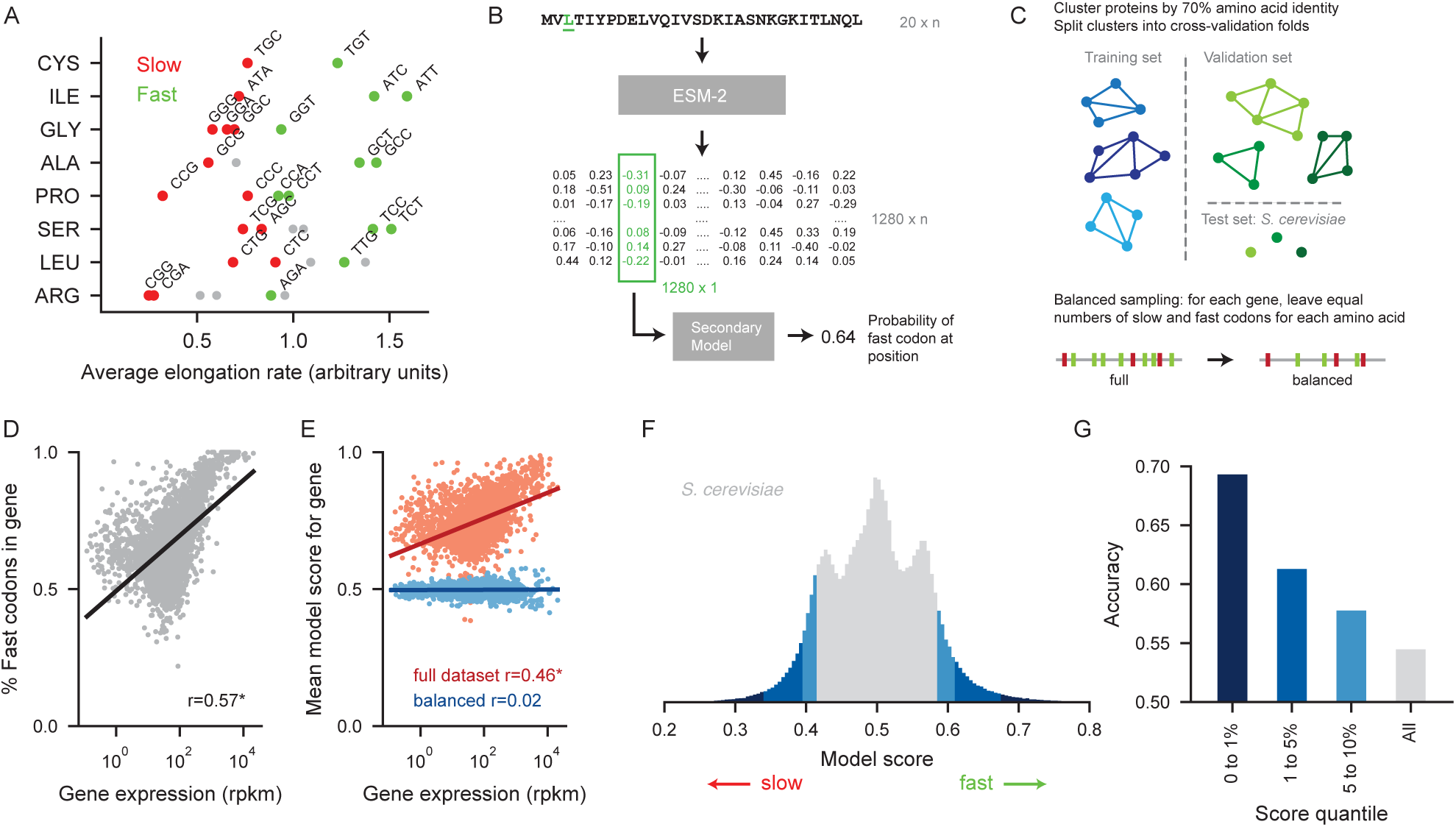
Model predicts codon choice solely from amino acid sequence. A) Codons were labeled as fast or slow based on the average per-codon elongation rate in *S. cerevisiae* as measured by ribosome profiling experiments. B) The entire amino acid sequence of a protein is fed into ESM-2, which outputs a representation of size 1280 x n, where n is the length of the protein. The representation vector at the position of interest is then fed into the secondary model, a neural network that outputs the probability of a fast codon at the position of interest. C) Proteins in the training dataset were clustered by 70% amino acid identity and clusters were split into five cross-validation folds. *S. cerevisiae* proteins were held out as a test dataset. For training and validation sets, positions were sampled to include equal numbers of slow and fast codons for each gene for each amino acid. In the test dataset, positions were weighted to place equal weight on slow and fast codons for each amino acid for each gene. D) The percentage of fast codons in a gene correlates strongly and significantly with the log of gene expression (Pearson’s *r* = 0.57). E) A model trained on the full dataset, without balancing, predicts more fast codons in genes with high expression (Pearson’s *r* = 0.46). We successfully removed this gene-specific effect by training on a balanced dataset with equal numbers of slow and fast codons for each amino acid for each gene (Pearson’s *r* = 0.02, not significant). F) Distribution of model scores for all positions in *S. cerevisiae*, where 0 indicates a slow codon and 1 indicates a fast codon. The model achieves an ROC AUC of 56.4%. The combined top and bottom 1%, 1-5%, and 5-10% quantiles are labeled in shades of blue. G) Accuracy for the combined top and bottom quantiles in the *S. cerevisiae* test dataset.

We then trained a neural network to predict whether each position of a protein should be encoded by a slow or a fast codon, built on top of Evolutionary Scale Model (ESM2) embeddings. ESM2 is a foundation model of protein sequence that learns a high dimensional embedding of each position for protein sequences from across the tree of life. ESM2 embeddings are created via unsupervised learning by training the model to predict what amino acid would be acceptable at masked positions of a sequence, and intrinsically reflect protein secondary structure, amino acid biochemistry, and other contextual information (Rives *et al*., 2021; Lin *et al*., 2023). Importantly, ESM2 is trained only on amino acid sequences and never observes the nucleotide sequence used to encode a protein. In our approach, we first extract a high-dimensional representation of a protein by feeding the entire amino acid sequence into ESM2. We then train a secondary neural network to predict the probability of a fast codon at a given position from the ESM2 representational vector at that position (Figure 1B).

Our model was trained on all available genes from thirteen non-*Saccharomyces* species of budding yeast and evaluated on *S. cerevisiae*. To prevent the model from simply memorizing conserved codons among homologous proteins, the proteins were clustered by 70% amino acid identity and the clusters were used to split the data into five training folds for cross-validation (Figure 1C). *S. cerevisiae* proteins were included in the clustering process but excluded from the training data and used as a test dataset. During evaluation, predictions were made on *S. cerevisiae* genes using the model that had not seen any proteins from the corresponding cluster during training. This approach of training the models only on non-*Saccharomyces* yeast and evaluating it on out-of-fold *S. cerevisiae* proteins forces the model to generalize further.

To compel the model to learn where in a given gene slow codons are more likely to occur, rather than the differences in overall codon composition between genes, we addressed several obvious sources of codon bias. Certain amino acids are more often encoded with fast codons, and in yeast, highly expressed genes use more fast codons than genes with low expression (Figure 1D). Although our model never sees expression data directly, a model that is aware of protein structure can learn to recognize that certain kinds of proteins are likely to have more fast codons. To remove these patterns, we balanced our training data by sampling equal numbers of slow and fast codons for each amino acid and each gene (Figure 1B). This sets the model’s baseline expectation that any particular position is encoded with a fast codon to be 50%, removing the correlation between model predictions and gene expression (Figure 1E).

Our model successfully identifies constraints on codon choice against a background of largely neutral positions. At the positions where the model output is closest to 0 (slow) or 1 (fast), indicating the highest certainty of the prediction, the model is notably accurate. It calls the correct label at 69.3% of the positions with the highest and lowest 1% of scores (Figure 1F,G). Its overall accuracy of 54.5% is robustly above the baseline accuracy of 50% for random guessing, demonstrating that meaningful information can be extracted from protein context even when codon choice at most positions is expected to be close to unconstrained. The extent of the constraint recoverable from protein information is equivalent to perfect predictability at 9% of positions, capturing the cumulative effect of small biases at a much larger number of positions.

### Cotranslational localization imposes constraints on codon choice

Having established that the model could learn constraints on codon choice, we next explored what biological contexts drive these constraints. We found a strong signal of slow codons at the start of genes. The model correctly predicts that slow codons are much more likely in the first 50 codons, with a mean model score of 0.465 (Figure 2A), compared to 0.503 after codon 50 (highly significant under Welch’s t-test with a moderate effect size of 0.54). We confirmed that this pattern was consistent for all amino acids, and thus not explained by differences in amino acid composition near the N terminus.

**Figure 2:**
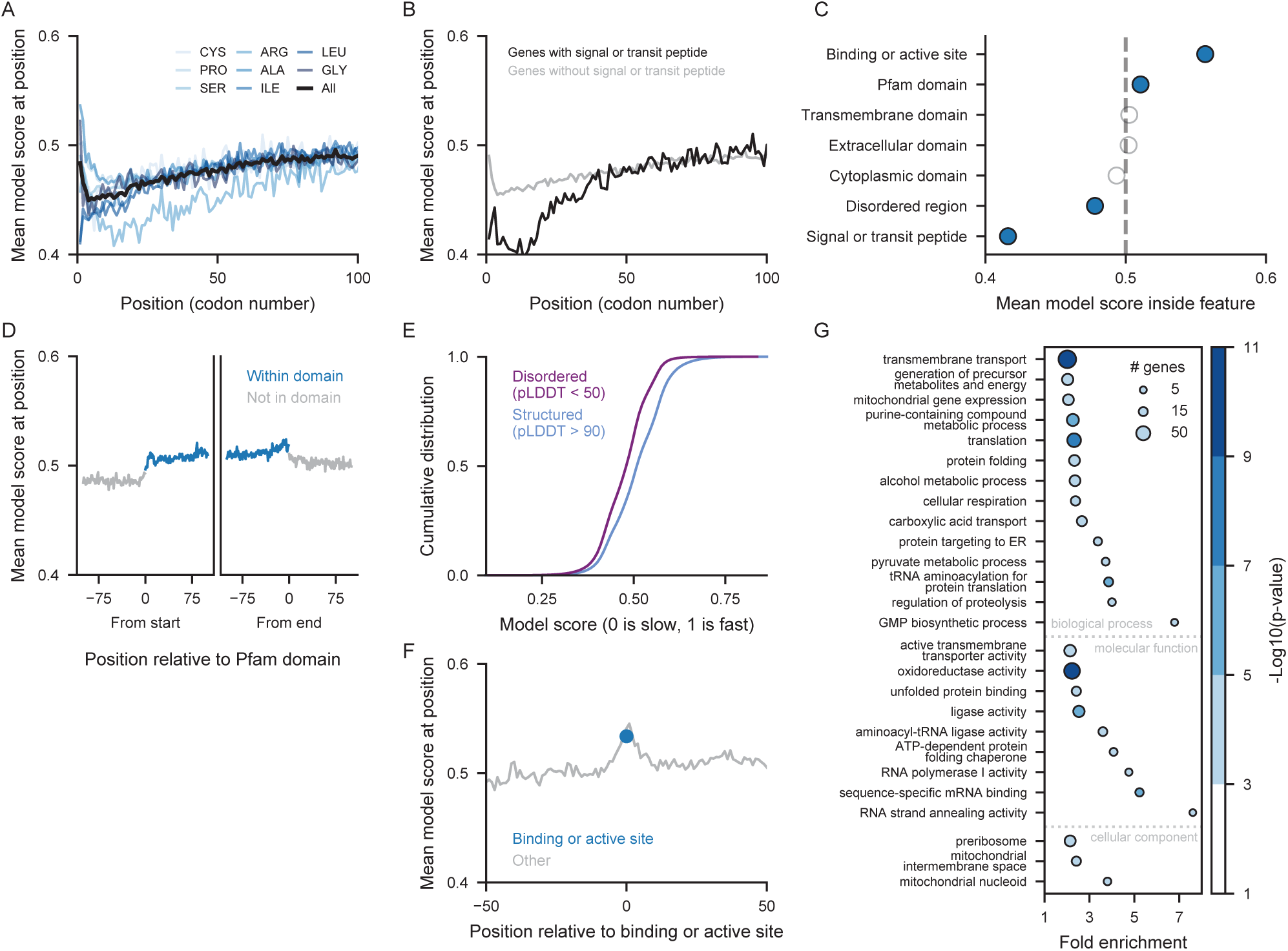
Model learns generalizable constraints on codon choice from position and protein structures. A) The model score at each position, averaged across all genes in the test dataset. Blue lines represent the mean model score at each position for positions with a specific amino acid. B) The model score at each position, averaged across all genes with a signal or transit peptide in black and across all genes without a signal or transit peptide in grey. C) The mean model score for positions inside the specified protein feature. Blue dots represent features with an effect size of at least 0.3. D) The mean model score at position relative to either the end or the start of a Pfam domain. Positions within a Pfam domain are plotted in blue and positions not inside of a Pfam domain are plotted in grey. E) The cumulative distribution of model scores inside of structured protein regions and disordered regions. The difference between the distributions is significant, with an effect size of 0.48. F) The mean model score at position relative to the start of a binding or active site. The mean model score of the first codon in a binding or active site is plotted in blue. G) GO terms enriched in genes with the lowest 10% mean model loss. Only GO terms with a false discovery rate < 0.05 and a fold enrichment > 2 are shown. Some overlapping and redundant categories were removed.

The preference for slow codons was particularly pronounced at the beginning of genes with signal peptides and mitochondrial transit peptides (Figure 2B). These peptides are located at the starts of proteins that must be localized to the ER or to the mitochondria, and encoding these peptides with slow codons might allow extra time for the signal recognition particle to bind to a signal peptide or for the mitochondrial transit peptide to be recognized by the TOM complex. In contrast to published reports (Pechmann *et al*., 2014), we do not observe a slowdown downstream of signal peptides; instead, the model predicts particularly slow codons at positions 5-15 (Figure 2B), well within the signal peptide. Inside of the signal peptide, the mean model score drops to 0.416 (Figure 2C). As above, this effect cannot be explained by differences in amino acid composition in signal and transit peptides (Figure S1). Instead, the model learns a true preference for slow codons within these important localization signals.

### Fast codons in structured regions and active sites suggest selection for translation accuracy

A long-standing hypothesis proposes that slow codons serve to separate individual protein domains during cotranslational protein folding. In this model, slow codons would allow the ribosome to pause when one domain of a multi-domain protein has emerged from the ribosome, allowing time for the domain to fold independently. Given the length of the ribosome exit tunnel, this would correspond to a region of slow codons 35-40 codons past the end of a domain. Previous studies have found some evidence that domains in prokaryotes such as *E. coli* are separated by stretches of slower codons (Zhang and Ignatova, 2009). The evidence in eukaryotes is less clear (Chaney *et al*., 2017). To test for this effect, we looked at codon usage around protein domains defined by Pfam. We do not observe any such slowdown 35-40 codons downstream of domain ends (Figure 2D).

Instead, we observe a preference for fast codons within domains and structured regions and slow codons in disordered regions (Figure 2C,D,E). Using AlphaFold’s pLDDT measure as a proxy for protein structure (Akdel *et al*., 2022), we find that structured regions (pLDDT > 90) have significantly higher model scores than disordered regions (pLDDT < 50), with an effect size of 0.48 (Figure 2E). We confirmed that this effect is not due to codon preferences at the beginning of genes; disordered regions at any position within a gene are predicted to have slower codons (Figure S2A). Nor is it due to a difference in amino acid composition, as the preference is still observed when amino acids are considered individually (Figure S2B).

The preference for fast codons within structured regions could arise from pressure for accurate translation. The ribosome samples tRNAs stochastically, in proportion to their availability, until it detects a correct tRNA with proper base pairing. Fast codons are matched by abundant tRNAs and can be decoded quickly, leaving less opportunity for accidental selection of the wrong tRNA and incorporation of the wrong amino acid (Mordret *et al*., 2019). Translation errors within structured regions have more extreme consequences, as amino acid substitutions disrupt structural interactions.

Active sites and ligand binding sites may be even more sensitive to amino acid changes and translation errors that change their functional interactions. Indeed, our model predicts a very strong preference for fast codons encoding these crucial amino acids. Positions within active sites and ligand binding sites have a mean model score of 0.557 (Figure 2F,C) corresponding to an effect size of 0.73 when compared to other positions within these genes. This effect cannot be explained just by the tendency of critical amino acid positions to fall within structured regions. Within structured regions, it is still true that active site and binding sites are predicted to have significantly faster codons, with an effect size of 0.48. Again, we confirmed that this preference for faster codons held true for amino acids considered individually. Our results highlight translation accuracy as a key driver of codon choice.

The associations between codon choice and amino acid identity, protein structure, and position are notable, but they combine to explain less than half of the total predictive power of our model. We quantified this by training a random forest to predict codon choice from just these three features, reaching a cross-validation accuracy of 51.8% on the test set. This suggests that only 40% of the gain in our model accuracy (54.5%) over baseline can be explained by simple and generalizable patterns of slow codons at the starts of genes and fast codons in structured regions. Our model must have captured more complex constraints on codon choice, such as patterns specific to particular protein structures or categories of genes.

### Codon constraints on transmembrane proteins and proteins localized to mitochondria

We next asked if some functional categories of genes had codon preferences that were captured particularly well by the model. We performed a gene ontology enrichment analysis on the 10% of genes with the best correspondence of true and predicted codon speed, as measured by the mean model loss across all positions in a gene. We found that this set of genes with accurate predictions was enriched for genes involved in transmembrane transport, genes localizing to the mitochondrial nucleoid and mitochondrial intermembrane space, and genes involved in protein folding (Figure 2G). These stronger predictions indicate specific biological constraints. Notably, the presence of proteins that function in membranes or inside the mitochondria aligns with our earlier observation of slow codons within localization signals. These genes may have particularly strong constraints on codon choice and translation speed to facilitate cotranslational localization. Chaperones such as heat shock proteins were also enriched among genes with low model loss. We hypothesize that it might be particularly important to facilitate their folding via translation speed, as proteins expressed in times of stress may need to be able to fold on their own without additional assistance.

### A competitive growth experiment identifies synonymous substitutions with fitness effects

Our model learns generalizable constraints on codon choice from evolutionary signatures that may have arisen from very weak selective pressure over a very long time. In many cases, these pressures would not be detectable in a short-term laboratory experiment, but we set out to find the important cases where codon choice is critical enough to create measurable phenotypes.

To map the landscape of fitness effects arising from synonymous mutations across the genome, we used high efficiency, precision genome editing to introduce thousands of synonymous mutations in endogenous yeast loci and then measured the resulting growth phenotypes. We used CRISPEY, a Cas9 retron editing system that enables extremely efficient sequence modification by homologous recombination of a donor DNA with the target locus after Cas9 cleavage (Figure 3A) (Sharon *et al*., 2018). We chose thousands of slow-to-fast and fast-to-slow synonymous substitutions to cover a wide range of model predictions, including positions where the model was confident in its selection of a fast or slow codon and positions where it did not show a strong preference. For amino acids with several codons with similar elongation rates, we also included synonymous mutations that would not alter elongation speed (slow-to-slow or fast-to-fast mutations). We constructed a pooled plasmid library with Cas9 guides and donor templates for these target substitutions.

**Figure 3:**
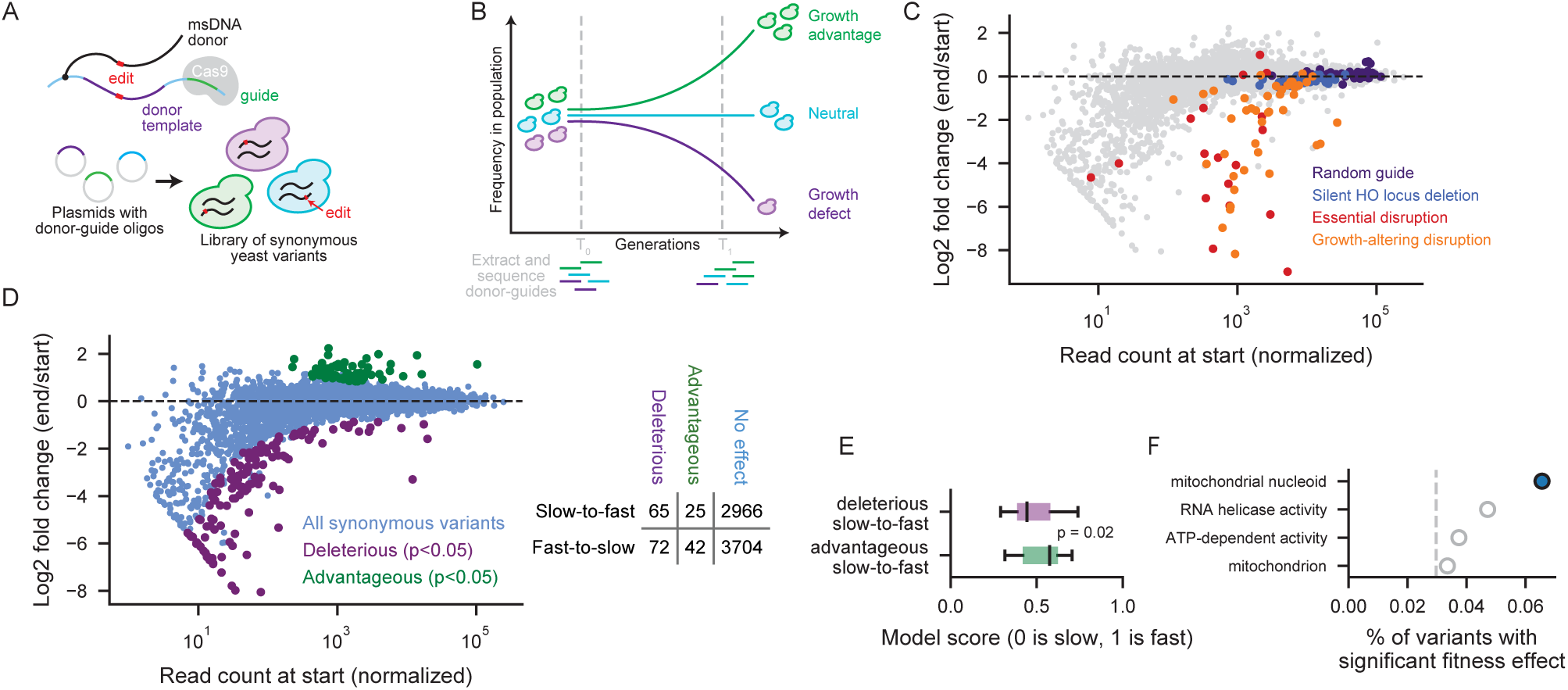
Genome-wide screen reliably identifies advantageous and deleterious synonymous mutations. A) Donor-guide oligos integrated into a plasmid backbone result in msDNA donor templates directly tethered to a Cas9 guide sequence, leading to efficient homologous recombination of edited sequences. This method was used to construct a yeast library with thousands of synonymous variants. B) Synonymous yeast variants were grown in competition for 81 generations. Plasmids carrying the donor-guide sequences were extracted from the population at the start and end of the competition. The variable region of the plasmids was sequenced to determine the change in population frequency of synonymous yeast variants at the start and end of the pooled competition. C) Change in frequency of control donor-guides from the start to the end of the pooled competition. D) Change in variant frequency from the start to the end of the pooled competition. Variants with significant effects on fitness were identified using DESeq2. E) Slow positions where a slow-to-fast mutation was deleterious had significantly lower model scores, indicating a preference for slow codons, compared to slow positions where a slow-to-fast mutation was advantageous. F) Four GO terms were found to be enriched among genes with synonymous variants with fitness effects, at a false discovery rate of < 0.1 and a fold enrichment > 1.5. To account for the variable number of positions tested in different genes, we measured the proportion of synonymous variants with a fitness effect among the genes associated with each GO term. We compared this to the overall percent of synonymous variants that have a significant effect on fitness (3%, indicated with dashed line). Blue represents a chi-squared p-value of less than 0.05.

We then measured the growth phenotype of each substitution in a pooled competitive growth assay. We grew our collection of yeast containing thousands of synonymous mutations in a pooled competition for 81 generations, with two competition replicates for each of two editing replicates, and collected samples at the start and end of the competition. The high efficiency of editing (94-96%) allowed us to use plasmid presence as a proxy for mutation presence. By sequencing the donor-guide region of the plasmids in each sample, we determined which synonymous variants were increasing or decreasing as a fraction of population over time (Figure 3B). To account for the overdispersion of sequencing data, we used DESeq2, a differential gene expression method, to determine which variants had a significant advantageous or deleterious effect on growth.

We first confirmed that our approach reliably distinguished true fitness effects from stochastic fluctuations. 59 of 60 negative controls showed no effect; these included guides targeting the inactive HO locus and non-targeting guides. The positive controls showed a deleterious effect for 14 out of 18 essential gene disruptions and 27 out of 44 disruptions of genes with known growth defects (Figure 3C).

Against this backdrop of robust experimental validation, our competition assay was able to identify a small but meaningful subset of synonymous mutations with measurable fitness effects, while most synonymous sites behaved neutrally. Among 6,874 distinct slow-to-fast and fast-to-slow mutations tested, we identified 3% with significant fitness effects: 137 deleterious and 67 advantageous (Figure 3D).

To validate results from our pooled sequencing-based approach, we conducted individual head-to-head growth competitions on five slow-to-fast synonymous variants identified by pooled competition. Individual variants, labeled with yellow fluorescent protein, were grown together with wild-type yeast labeled with red fluorescent protein. The proportion of variant cells over time, measured over four days, provided a direct measure of fitness. The fitness effect was reproducible for four out of the five variants (Figure S3).

We note that our results stand in contrast to a recent report that the majority of synonymous mutations in yeast affect cell fitness. We attribute this discrepancy to key differences in the statistical model for detecting a deviation from the null hypothesis; our controls show that our approach successfully limits false discoveries. Three synonymous yeast variants that Shen *et al*. measured as deleterious were serendipitously present in our pooled library, and none of these variants affected yeast fitness in the conditions tested (synthetic media as opposed to the rich media used in Shen *et al*.).

Our results show that most synonymous substitutions have no measurable effect in laboratory growth conditions, matching long-standing expectations that most such changes are nearly silent. Against this background, we reveal a small set of synonymous changes with a robust fitness effect detectable over the course of a short competitive growth experiment.

### Fitness effects of mutations correspond to model predictions

Although our deep learning model focused specifically on decoding speed, our competitive growth experiment was agnostic to the underlying mechanism by which a synonymous substitution affected fitness. However, for slow positions, we do observe a correspondence between model predictions and experimental results. Among positions where a mutation from a slow to a fast codon caused a growth defect, the model was more likely to predict a slow codon (median score of 0.44), and among positions where a mutation from a slow to a fast codon caused a growth advantage, the model was more likely to predict a fast codon (median score of 0.58; significantly different with p = 0.021 from a Wilcoxon rank-sum test) (Figure 3E). We do not see such correspondence for fast positions.

The experiment also detects fitness effects that do not relate to translation speed. For amino acids with several codons with very similar elongation rates, our library included mutations from one slow codon to another or from one fast codon to another. We found that, overall, synonymous mutations that change the translation speed were no more likely to cause significant effects on yeast fitness than synonymous mutations with no change in translation speed. Our model and our experiment do find synonymous mutations that disrupt biology by changing translation speed, but the experiment also finds many positions where the nucleotide sequence itself matters independent of the decoding speed of the codon. The many positions where synonymous changes have fitness effects reflect the wide range of post-transcriptional processes that rely on mRNA sequence.

### Genes localizing to the mitochondrial nucleoid are sensitive to synonymous mutations

Fitness-altering synonymous mutations disproportionately affected proteins targeted to the mitochondria, showing a remarkable correspondence with the predictions of our model (Figure 3F). After accounting for the number of positions tested in each gene, proteins that localize to the mitochondrial nucleoid were the sole overrepresented GO category among genes with fitness-altering mutations. Indeed, we found synonymous mutations with fitness effects in 6 out of the 7 genes in this category that were present in the experiment, representing 8 of 122 positions tested in these genes. Interestingly, this is not solely due to codon preferences within localization signals; none of these mutations fell within transit peptides at the start of the gene. The necessity of importing

## Discussion

Translation is the central function of the genetic code, and this process imposes selective pressures on the choice between synonymous – yet not identical – codons. These pressures intertwine with pressures from many other aspects of gene expression to shape the coding sequence of a gene. Our study cuts through these other forces to reveal biological constraints imposed on codon choice by the process of translation itself (Figure 4).

**Figure 4:**
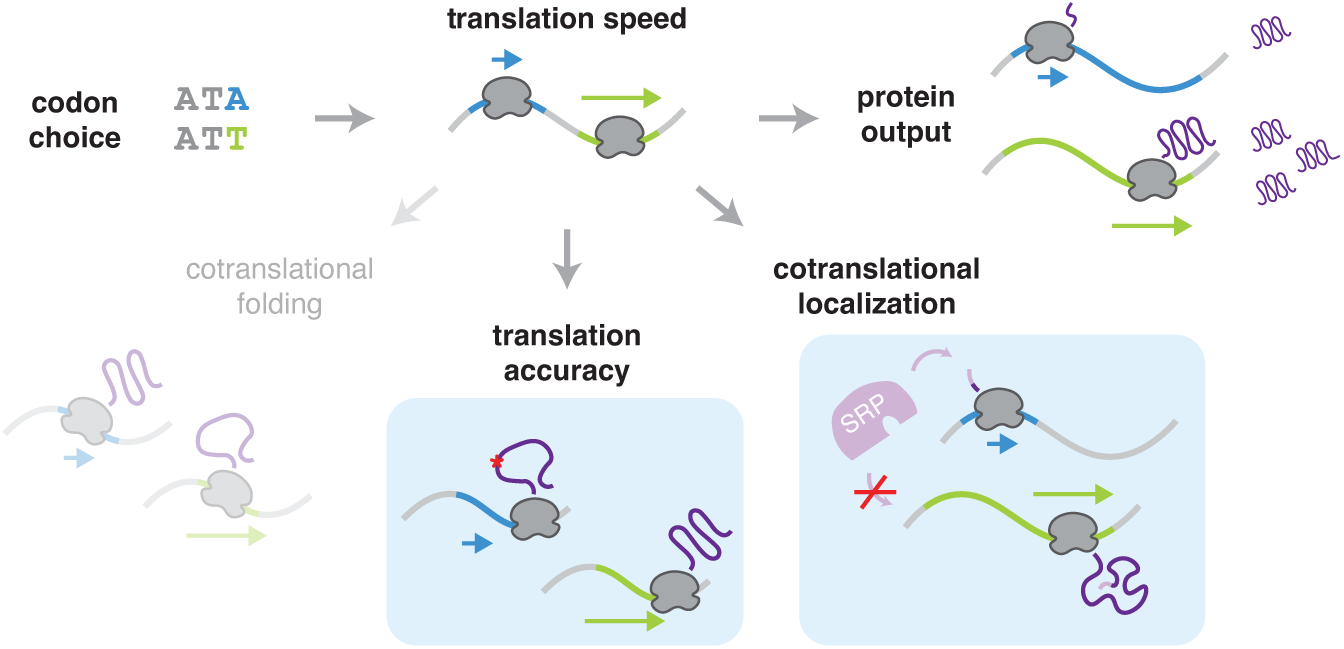
Codon choice affects translational accuracy and cotranslational localization. Synonymous mutations affect the translation speed of a ribosome. Speed can directly impact how much protein is produced from an RNA. Our study reveals that codon choice in eukaryotes is also constrained by translation accuracy, with a preference for fast codons in important structured regions, and by cotranslational localization, with a preference for slow codons near the beginning of some transcripts to provide extra time for proteins such as SRP to bind to the nascent protein. We find a limited impact of constraints from cotranslational folding in eukaryotes. these proteins across two membranes in the mitochondria may impose particularly strong constraints on their codon usage and nucleotide sequence. Both experiment and model suggest that cotranslational localization is a dominant source of position-specific constraints on codon choice.

Our deep learning model implicitly synthesizes evolutionary information to identify generalizable, subtle constraints on codon choice. Our work goes beyond the well-characterized preference for fast codons in highly expressed genes to find individual positions within specific genes where codon choice seems constrained. By using a model built from protein sequence alone, and by forcing the model to ignore differences in codon composition between genes, we specifically probe the extent to which protein structure and function influence codon choice. By constructing a training set that deals thoughtfully with homologs, we prevent our model from simply memorizing gene-specific codon patterns that may not generalize across genes. Through this careful approach, we are able to predict constrained positions where synonymous mutations have small but seemingly genuine influence, visible over the course of evolution but unlikely to be detectable in a short-term growth experiment.

Our results show that it is in fact possible to uncover real biological signal relating to codon choice from a deep evolutionary model of protein sequence alone. Our model recovers constraints equivalent to perfect prediction at 1 in 11 positions, capturing the cumulative effect of small biases at a much larger number of positions. We are able to find signs of preference for fast codons within structured regions and especially at interaction sites, perhaps arising from a greater need for translation accuracy at these functionally important positions, as slow decoding of non-preferred codons would allow more chances for harmful amino acid misincorporation. Conversely, we find a preference for slow codons near the beginnings of genes and especially in signal and transit peptide sequences, perhaps reflecting the importance of slow translation to allow proper cotranslational localization. Counter to results in bacteria, we do not find strong signatures of cotranslational folding, with no clear trend of slow codons downstream of independently folded protein domains.

In contrast to the subtle but general constraints learned by our model, our competitive growth experiment captures positions where a single codon is critical to a specific protein. We reliably identified a set of about two hundred positions at which synonymous changes caused a noticeable effect on cell fitness, while the vast majority of synonymous changes did not measurably affect growth. These critical positions may each be highly specific to an individual protein and not representative of general trends. The mechanisms by which these synonymous mutations impact cell fitness are not necessarily related to translation. Rather, they may arise from nucleotide constraints at many different levels, from transcription to mRNA structure and half life. Indeed, synonymous mutations that do not change the decoding speed were as likely to show effects as those that do change the speed, implicating constraints unrelated to translation. These mutations thus highlight the value of our evolutionary analysis that can find subtle constraints likely to arise directly from the process of translation.

In light of the different information captured by our two approaches, it is particularly noteworthy that common trends emerge. Both approaches suggest that genes encoding proteins localizing to the inner mitochondrial space have particularly strong constraints on codon choice. These proteins must be imported past both the outer and inner membrane of the mitochondria, and previous work has shown that slow codons, either within transit peptide sequences or elsewhere in the gene (Tsuboi *et al*., 2020), can be important for their localization.

Codon usage bias has been a longstanding topic of interest in molecular and evolutionary biology, and modern applications like synthetic mRNA vaccines have highlighted the importance of more sophisticated approaches to codon optimization. We have shown that the nuanced rules of mRNA sequence go beyond simple preferences for ‘optimal’ codons. Our results demonstrate that protein structure predictably constrains codon choice in ways that can be recognized by a model that only considers amino acid sequence. The ability to predict codon choice from protein sequence is a step towards rational mRNA design and an improved understanding of pathogenic synonymous mutations.

## Data availability

Analysis code and data are available at https://github.com/lareaulab/codon_constraints and raw sequencing data are available at the NCBI Sequence Read Archive via accession number PRJNA1297196.

## Acknowledgments

This work was supported by the National Institutes of Health (National Institute of General Medical Sciences R01GM132104 to L.F.L.), the National Science Foundation (1936069 to L.F.L.), the Rose Hills Foundation (to L.F.L.), the Shurl and Kay Curci Foundation (to L.F.L.), and the Chan Zuckerberg Biohub (to L.F.L.). H.S. was supported by the Department of Defense through the National Defense Science & Engineering Graduate Fellowship (NDSEG) Program. We thank the Vincent J. Coates Genomics Sequencing Laboratory (QB3 Genomics, UC Berkeley, Berkeley, CA, RRID:SCR_022170; supported by NIH S10 OD018174 Instrumentation Grant) and the Chan Zuckerberg Biohub for sequencing support, and the U.C. Berkeley flow cytometry core facilities. We thank Nicholas Ingolia for thoughtful conversations.

## Methods

### Definition of fast and slow codons

Codons were labeled in a binary manner as either fast or slow based on empirical elongation rates from ribosome profiling experiments, computed as the inverse of the average ribosome footprint density for each codon (*e.g.*, across all ‘CGA’ codons) by Weinberg *et al*. (2016). We limited our consideration to eight amino acids with a difference of at least 0.35 in the elongation rate of the slowest and fastest synonymous codons: cysteine, proline, serine, arginine, alanine, leucine, isoleucine, and glycine. Codons with an elongation rate less than 0.85 were labeled as slow, and codons with an elongation rate greater than 0.85 were labeled as fast (except that for leucine, CTC was labeled as slow with an elongation rate of 0.91). We excluded any codons with an intermediate elongation rate.

Codon preference and elongation rate can vary between species, potentially confounding analysis. To consider only codons where the translation rate does not vary between species, we calculated tAI scores from tRNA gene counts according to dos Reis *et al*. (2004). We obtained tRNA gene counts from GtRNAdb (Lowe and Chan, 2016) on September 6, 2023. For *S. cerevisiae*, we used tAI measurements provided in Weinberg *et al*. (2016). If tAI scores for a codon were variable between species, implying that the translation rate of that codon might change greatly from species to species, the codon was dropped from consideration. Thus, the set of codons considered by the model can be assumed to be translated consistently slow or consistently fast among the different yeast species in the training set. After this process, 14 slow codons and 12 fast codons belonging to eight amino acids remained in consideration (Figure 1A).

### Training dataset

The training dataset consisted of amino acid sequences of all proteins from thirteen species of non-*Saccharomyces* budding yeast (order *Saccharomycetales*). Genomic coding sequences and protein sequences were downloaded from NCBI Datasets on September 5, 2023, and coding sequence with no corresponding protein sequence in the dataset were removed. To limit analysis to proteins with homologs in multiple species, all proteins were clustered using MMseqs2 (Steinegger and Söding, 2017) at an amino acid identity of 40% and clusters with fewer than four sequences were removed. The final training set comprised 45,274 proteins.

The sequences were then split into five training folds for cross-validation. To ensure that sequences in a training fold were not homologous to sequences in the validation fold, training sequences as well as *S. cerevisiae* test sequences were clustered by 70% amino acid identity using MMseqs2. All training sequences in a cluster were then randomly assigned to the same training fold. We used each amino acid sequence as an input to the 650M-parameter ESM2 model (Lin *et al*., 2023), and retrieved a high-dimensional representation of 1280 features per position in the sequence. We use these per-position representations as inputs to our model. ESM2 has a maximum context length of 1024, so longer amino acid sequences were truncated. We randomly undersampled our training dataset to leave an equal number of slow and fast codons per amino acid for each gene, resulting in a total training set size of 3.5 million positions.

### Model

Our model consists of a four-layer deeply connected feed-forward neural network with two layers of size 128 and two layers of size 64. The model has a total of 192961 trainable parameters. The layers use ReLu activation, and binary cross entropy was used as the loss function. The model was trained with the Adam optimizer (learning rate 0.0001), a batch size of 128 and a dropout rate of 50%.

### Model evaluation and analysis

The model was evaluated on all *S. cerevisiae* protein sequences (S288C, R64-1-1), excluding dubious sequences and retrotransposons. Sequences and annotations were obtained from the UCSC Genome Browser on July 8, 2020. To ensure that the model was forced to learn generalizable information, predictions on test sequences were made using a model that had not seen any homologous sequences during training. Specifically, for each test sequence, predictions were generated using the particular cross-validation model that was not trained on any sequences that clustered with the test sequence at 70% amino acid identity. Note that no *S. cerevisiae* sequences were used during model training, enforcing an additional layer of generalization. To maintain the same class balance used in the training regime, positions in the test dataset were weighted to place equal weight on fast and slow positions for each gene for each amino acid. Weights were normalized so that the total weight for each amino acid for each gene was equal to twice the number of positions with the underrepresented class.

When comparing the weighted mean accuracy between genes of low and high expression, gene expression measurements were obtained from Weinberg *et al*. (2016). Protein feature annotations were obtained from Uniprot on September 26, 2023 and Pfam domain annotations were obtained from SGD on October 7, 2024. When analyzing the connection between model scores and protein structure, we used pre-computed pLDDT scores obtained from the AlphaFold Protein Structure Database (Varadi *et al*., 2024; Jumper *et al*., 2021) on April 7, 2025. Statistical significance of all comparisons was assessed using a weighted variant of Welch’s t-test, and effect sizes are reported in terms of Cohen’s *d*.

### Random forest comparison

We trained a random forest model using pLDDT, position, and amino acid identity as input. The accuracy of the random forest model was assessed as the mean accuracy across 5 cross-validation folds on the *S. cerevisiae* test set. Hyper-parameters were tuned via grid search, and the final parameters used were 500 trees with a maximum depth of 10, with at least 5000 samples required to split a node.

### Synonymous variant design

For our genome-wide screen for synonymous mutations with effects on cell fitness, we chose to target 3068 unique slow-to-fast and 3818 unique fast-to-slow mutations across 1238 different genes. Target mutations were chosen to represent positions with a range of model scores, including both positions the model considered likely to be slow or likely to be fast, and positions that had intermediate model scores. Target positions were chosen to include a comparable number of slow and fast codons from each amino acid. All genes targeted were either essential or known to cause at least a 10% growth rate defect when knocked down (Breslow *et al*., 2008). Some categories of genes, such as genes involved in chromosome separation, DNA repair, DNA replication, and plasmid maintenance, were also excluded from the study because of our reliance on plasmid counts to read out mutation effects.

For each target synonymous mutation, we designed a donor-guide oligonucleotide as specified by Sharon *et al*. The 194 bp oligonucleotide consists of a 5’ homology region, a unique 100-bp long donor sequence carrying the desired single-codon edit, a retron sequence, a 20-bp long Cas9 guide sequence, and a 3’ homology region. We chose guide sequences that would not have any off-target effects by aligning all candidate guides against the entire yeast genome with bowtie. If a guide aligned to any off-target region with a valid PAM site and fewer than three mismatches, it was excluded. We also ensured that the desired mutation would change the genomic DNA to prevent repetitive cutting by Cas9, either by disrupting the seed region (first seven nucleotides from the 3’ end) or PAM site used by the guide. Lastly, we removed any donor-guide oligonucleotide that contained a repeat of length 10 or longer, due to difficulty of synthesis. For 2007 of our target mutations, more than one guide fulfilled all the criteria, and a second donor-guide targeting the same mutation was included.

For certain amino acids that had several codons with very similar elongation rates, we also included slow-to-slow and fast-to-fast mutations alongside our slow-to-fast and fast-to-slow mutations. These mutations include mutations between the slow codons CGA and CGG (arginine), GGA and GGC (glycine), TCG and AGC (serine), and between the fast codons GCC and GCT (alanine), CCT and CCA (proline), and TCT and TCC (serine). We included complementary slow-to-slow or fast-to-fast mutations for 2215 of our slow-to-fast or fast-to-slow target mutations.

Lastly, we included several sets of donor-guide oligos as controls. We included 42 random guide sequences that do not align anywhere in the yeast genome. We also target 33 random 2-base pair deletions to the HO mating locus, where any changes should not affect the fitness of haploid yeast. Lastly, we also include a set of controls intended to knock out 57 different essential genes or genes with known growth defects. These control donor-guides introduce ATGN-to-TAA mutations designed to both remove potential start codons, and, if there is more than one start codon for the gene, to introduce a premature stop codon and frameshift. In total, we designed a library of 12,000 donor-guide oligos, which was then produced by Twist Biosciences.

### Synonymous yeast library construction

The donor-guide oligonucleotide library was assembled together with the pZS165 background as described in Sharon *et al*. The assembled plasmid library was cleaned and concentrated using DNA Clean & Concentrator-5 from Zymo Research. The assembled plasmids were electroporated into Endura electrocompetent *E. coli* cells in 1 mm cuvettes using electroporation settings of 1800 V, 200 Ω, and 25 *µ*F. Two electroporation reactions were performed, each using 1 *µ*L of 36 ng/*µ*L assembled plasmids. The transformed bacteria were incubated in the provided recovery medium at 37 C, 300 rpm, for 1 hour. Serial dilutions were plated in order to evaluate transformation efficiency, and remaining cells were incubated at room temperature in 200 mL of LB + Carbenicillin overnight at 220 rpm. Once the target transformation efficiency (1000 transformed cells per donor-guide) was confirmed via dilution plates, bacteria were incubated at 37 C to an OD of 1.5 to 2. A Qiagen HiSpeed Midiprep kit was used to harvest assembled plasmids from the bacteria. The plasmid library was then digested with Not1 and CIP to remove empty vectors, cleaned, and concentrated.

A background yeast strain was constructed by integrating a galactose-inducible Cas9 and retron transcriptase into BY4742, as specified in Sharon *et al*., 2012. A scaled chemical yeast transformation was then performed (Gietz and Schiestl, 2007). After plating serial dilutions, remaining yeast cells were incubated overnight in 500 ml of SD -HIS/-URA + dextrose at 30 C, and an initial OD measurement was taken. Once the yeast reached an OD of 0.5 to 0.75 (after subtracting the initial OD measurement), the yeast were spun down for 5 minutes at 4000 and frozen at -80 C in 15% glycerol stocks. A total of 1.5 million transformants were generated (more than 1000 transformed yeast cells per donor-guide).

### Pooled genomic editing

To induce genomic editing, we performed two separate editing replicates. We started the pre-editing transformed yeast libraries in 250 mL of SD -URA/-HIS + Raffinose at an OD of 0.12. After 24 hours, yeast cultures were spun down at 3000 rpm, and transferred to 250 mL of SD -URA/-HIS + Galactose at an OD of 0.1 to 0.3. This step was repeated for a total of 48 hours of growth in galactose media.

After editing, yeast were spun down and transferred into 250 mL of SD -URA/-HIS + Dextrose at a starting OD of 0.12, and allowed to recover for six hours before the start of the pooled competition. All incubation steps were performed at 30 C and 220 rpm. Editing efficiency was validated by Sanger sequencing on edited colonies from a set of 8 different donor-guides, and successful HDR was confirmed for 59 out of 63 colonies. The observed editing efficiency of 94% is close to the reported 96% efficiency (Sharon *et al*., 2018).

### Pooled competition

At the start of the competition, each of the two editing replicates was split into two experimental replicates for a total of four experimental replicates. Samples were taken from each replicate, pelleted at 3000 rpm, and frozen at -80 as a pellet with the supernatant removed. Yeast were then cultured together over the course of 12 days. Yeast cultures were transferred to new media every day, and the OD was allowed to vary from 0.05 to 2.8. Prior to taking a sample, yeast were back-diluted to an OD of 0.1 and allowed to grow for 6 hours to an OD of 0.8. The final sample was taken after 81 generations.

To retrieve plasmids carrying donor-guide sequences from the frozen yeast pellets, we used a modified protocol (Muller *et al*., 2022) for the Zymo Yeast Miniprep II kit. Each frozen yeast pellet contained a total of 30 OD units of yeast, and was resuspended in 800 *µ*L of solution 1 with 30 *µ*L of zymolyase, and digested for 3 hours at 37 C, 900 rpm. Scaled amounts of solution 2 (400 *µ*L) and solution 3 (800 *µ*L) were added as specified by the kit. Two centrifugation steps were then performed at 2.1k g for 10 minutes, transferring the supernatant each time. The resulting supernatant was then sequentially passed across the same column, and DNA was eluted from the column in 30 *µ*L of 37 C water after 5 minutes of incubation at 37 C. The resulting DNA was further cleaned and concentrated.

### Sequencing and analysis

Donor-guide sequences were amplified from the retrieved plasmid library using primers 5 and 6. Primer 6 is composed of 8 separate primers of varying length that add 1 to 8 extra nucleotides to increase library diversity for sequencing. The amplification was conducted using Q5 high-fidelity polymerase (New England Biolabs) for 12 PCR cycles with a melting temperature of 63 C. Samples were cleaned and concentrated before adding dual-indexed NEBNext Multiplex Adaptors. Adaptors were added using Q5 polymerase for 7 PCR cycles with a melting temperature of 70 C. All eight samples (start and end for four competition replicates) were then run together on one lane of NovaSeq X as 100bp single-end reads according to the manufacturer’s protocol by the Vincent J. Coates Genomics Sequencing Laboratory at the University of California, Berkeley.

Reads were aligned to the donor library using bowtie with parameters ‘-n 2 -l 58 –best –trim5 28’. The first 28 bases of each read, corresponding to the 5’ vector homology sequence, were trimmed. Only reads aligning as expected to the first 72 bases of a donor sequence were counted. Each sample contained 60 - 150 million reads with a valid alignment. To call synonymous variants with significant changes to population frequency between the start and end of the competition, we used DESeq2, a differential gene expression method which accounts for the relation of measurement variance to the mean, and normalizes read abundances across samples. We use custom normalization factors estimated from the top 75% most abundant reads. Variants with significant fitness effects were called based on a Wald test of the absolute value of the log2 fold change being greater than 0.58, which corresponds to at least a 0.72% change in growth rate compared to wild-type. A variant was considered to have a significant fitness effect if the adjusted p-value (alpha = 0.01) was less than 0.05.

### GO Analyses

GO enrichment analyses were conducted using the PANTHER Overrepresentation Test (released 2024-08-07) using GO Ontology database annotations (released 2024-11-03, DOI: 10.5281/ zenodo.14083199). We used a Fisher’s exact test with FDR correction, and performed GO enrichment analyses on the GO biological process complete, GO molecular function complete, and GO cellular component complete annotation sets. When testing for enrichment among low-model-loss genes, the reference list was composed of all genes in the test set with a total weight of at least 20 (generally, genes with at least 10 slow positions). The analyzed list was composed of the genes with model loss in the lowest 10% among the reference genes. When testing for enrichment among genes with significant fitness effects in the pooled competition, the reference list was composed of all the genes with fast-to-slow or slow-to-fast variants included in the yeast library. The analyzed list was composed of genes that had at least one fast-to-slow or slow-to-fast mutation with a significant effect on fitness as measured by the pooled competition.

### Head-to-head competition

We chose five synonymous variants to test individually in order to confirm the reliability of the pooled competition via paired fluorescence-labeled competition. The synonymous variants were chosen from a previous pooled growth competition in which randomly-chosen slow-to-fast variants were grown for 19 generations. An additional mutation, a two-nucleotide deletion to the start codon of CNN1, was known to have a small deleterious effect on cell fitness and was chosen as a positive control.

We constructed background yeast strains as described previously, with the addition of an integrated fluorescent protein. Using an EasyClone 2.0 integrative vector (Stovicek *et al*., 2015), a red (mCherry) or yellow (eCitrine) fluorescent protein sequence was integrated into a genomic locus and selected for using hygromycin. We then ordered the five corresponding donor-guide oligos, assembled the plasmids, and transformed them into yellow fluorescent yeast as described above. As a wild-type control, we transformed the empty backbone plasmid pZS165 into both the red and yellow fluorescent yeast.

After confirming the presence of the desired plasmid, Cas9-mediated editing was induced in all 9 strains by growing the yeast in SCD -URA/-HIS + Raffinose for 24 hours, then in SCD -URA/-HIS + Galactose for 48 hours, and lastly in SCD -URA/-HIS + Dextrose for an additional 6 hours. Cells were grown in a block in 2 ml volumes at 37 C and 750 rpm. After editing, cells were spread on SCD -URA/-HIS plates. Colonies were Sanger-sequenced to confirm the presence of the desired mutations at the corresponding genomic loci, and three editing replicates were chosen for each variant.

For the head-to-head competition, yellow-fluorescent variant yeast were mixed with red-fluorescent wild-type yeast at a ratio of 1 to 1 at an OD of 0.04 and allowed to grow for 18 hours. After this, they were diluted with a ratio of 1-in-10, and allowed to grow for 6 hours to an OD of 0.5 before a 200 *µ*L sample was taken and fixed with 4% PFA. Yeast were then diluted at a ratio of 1-in-20 and allowed to grow for another 18 hours, to an OD of 1. This cycle was repeated four more times for a total of five timepoints, allowing the variant yeast to grow in competition with the wild-type yeast for approximately 30 generations.

Sampled cells were pelleted by five minutes of centrifugation at 3000 rpm, and washed once with DPBS before being resuspended in 200 *µ*L of 4% PFA and incubated for 30 minutes in the dark at room temperature. Fixed samples were then washed twice with DPBS and stored at 4 C for less than a week. Before analysis, samples were diluted 1-to-4 in DPBS. A BD LSRFortessa analyzer was used to count the number of yellow-fluorescent variant cells and red-fluorescent wild-type cells in each sample. For each fixed sample, fluorescence was measured for 10,000 cells, and cells were labeled as red or yellow with a custom script. The relative growth rate was calculated as described in Breslow *et al*. (Breslow *et al*., 2008).

## SUPPLEMENT

**Figure S1:**
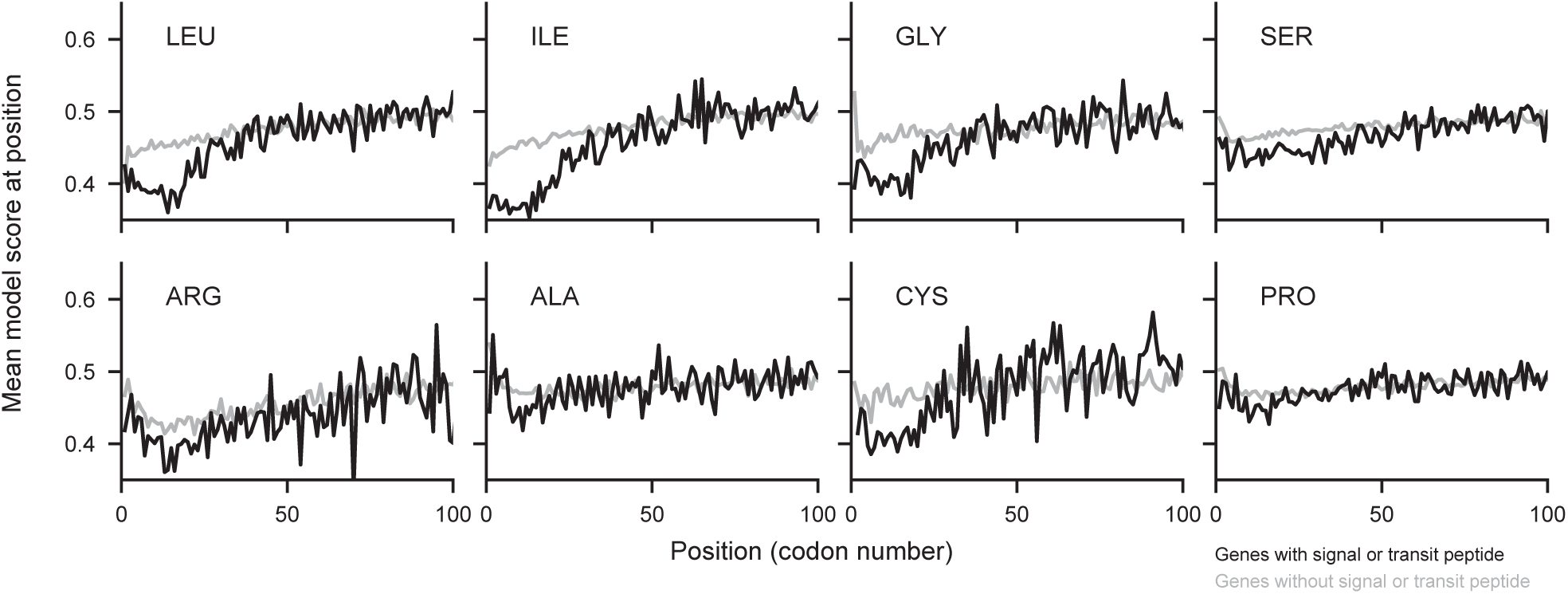
Signal and transit peptides are encoded with slower codons for all amino acids. The mean model score at each position with a given amino acid, averaged across all genes with a signal or transit peptide in black and across all genes without a signal or transit peptide in grey. For all amino acids, the first 50 codons are predicted to be significantly slower in genes with signal or transit peptides compared to other genes. The effect size is strongest for isoleucine (0.73) and leucine (0.64) and weakest for alanine (0.15) and proline (0.12).

**Figure S2:**
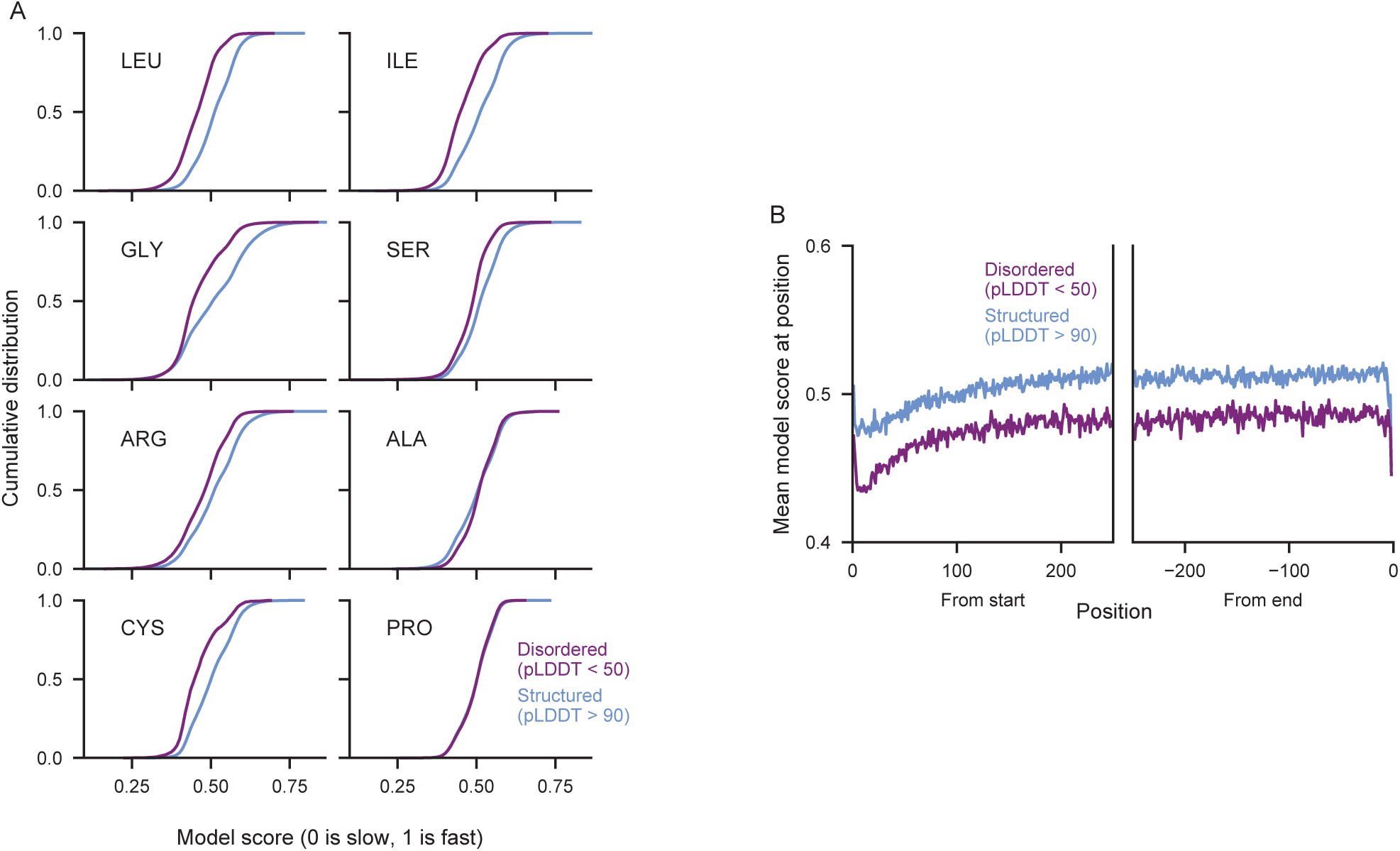
Structured regions predicted to be encoded with faster codons regardless of amino acid or position. A) The cumulative distribution of model scores for positions in structured (pLDDT > 90) and disordered (pLDDT < 50) regions, plotted individually for each amino acid. Positions in structured regions are predicted to have significantly more fast codons than positions in disordered regions for all amino acids except proline and alanine. B) The mean model score at positions in structured (blue) or disordered (purple) regions, plotted by position from the start and from the end of the gene.

**Figure S3:**
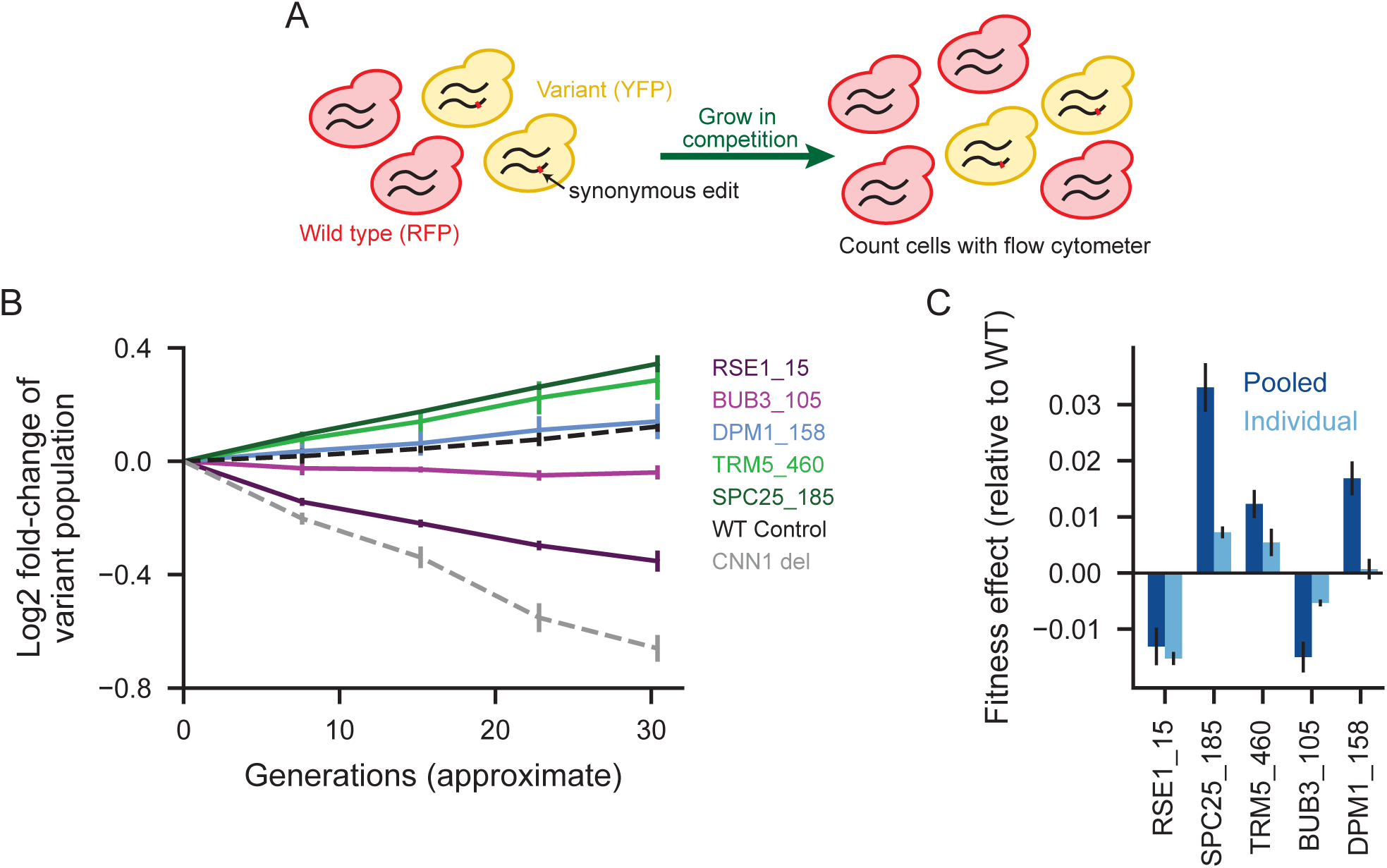
Paired fluorescently labeled competition. A) Yellow-fluorescent cells containing a specified synonymous slow-to-fast mutation were grown in competition with red-fluorescent wild-type cells. A flow cytometer was used to count the proportion of variant cells over time. B) The change in proportion of variant cells over thirty generations for five synonymous slow-to-fast mutations, a yellow-fluorescent wild-type control, and a gene deletion control known to have a weak growth defect. C) The measured fitness effects of the synonymous variants in the paired head-to-head competition compared to the fitness effects observed in a pooled competition.

